# Multi-laminate Annulus Fibrosus Repair Scaffold with an Interlamellar Matrix Enhances Impact Resistance, Prevents Herniation and Assists in Restoring Spinal Kinematics

**DOI:** 10.1101/418103

**Authors:** Ryan Borem, Allison Madeline, Ricardo Vela, Sanjitpal Gill, Jeremy Mercuri

**Author notes:** Corresponding Author Jeremy Mercuri, Ph.D. 313 Rhodes Engineering Research Center Clemson, SC, 29634 (email).

## Abstract

Focal defects in the annulus fibrosus (AF) of the intervertebral disc (IVD) from herniation or surgical injury have detrimental impacts on IVD mechanical function. Thus, biomaterial-based repair strategies, which can restore the mechanical integrity of the AF and support long-term tissue regeneration are needed. Accordingly, a collagen-based multi-laminate scaffold with an underlying “angle-ply” architecture has been previously reported demonstrating similar mechanical properties to native AF tissue. The objectives of this work were to: 1) enhance the biomaterials impact strength, 2) define its contribution to spinal kinematics, and 3) assess its ability to prevent IVD herniation. First, AFRP’s were enriched with a glycosaminoglycan-based (GAG) interlamellar matrix (ILM), and then tested for its radially-directed impact resistance under physiological stresses. Subsequent kinematic testing was conducted using a characterized GAG-enriched AFRP as an AF focal defect closure device. In summary, AFRPs demonstrated 1) incorporation of a GAG-based ILM significantly increased radial impact strength, 2) restoration of axial FSU kinematics and 3) ability to prevent herniation of native IVD tissues. Together, these results suggest that the AFRP demonstrates the mechanical robustness and material properties to restore an IVD’s physiological mechanical function through the adequate closure of an AF focal defect.

## 1. Introduction

Intervertebral discs (IVDs) consist of a gelatinous core known as the nucleus pulposus (NP), which is circumferentially constrained by a fiber-reinforced, multi-laminate structure known as the annulus fibrosus (AF). The NP’s hydrophilic extracellular matrix (ECM) is composed of aggrecan and collagen type II, while the AF is composed of 15-25 concentric lamellae of collagen type I with fibers aligned in a ± 28-43° “angle-ply” microarchitecture. (Urban and Roberts, 2003) Between each AF lamellae, non-fibrillar ground substance (e.g., glycosaminoglycan; GAG), elastin, collagens type I and VI, and cells form an inter-lamellar matrix (ILM). (Tavakoli et al., 2016) The AF’s highly organized, fiber-reinforced structure resists tension and torsion during spinal bending and twisting, but also bears hoop stresses developed from pressurization of the NP during compressive loading of the IVD. (Ducheyne, P. Healy, 2011; R. G. Long et al., 2016; Patwardhan et al., 2003; Schultz et al., 1982) Furthermore, the ILM reinforces the AF against these radially-directed intradiscal pressures (IDPs) and prevents de-lamination of adjacent lamellae. (Adam et al., 2015) Thus, the ECM composition and organization of these IVD tissues allow for stable, multi-axial motion of the spine through their mechanical interplay.

Physiologically, human lumbar IVDs experience axial compressive loads that generate IDPs ranging from 0.1-2.3 MPa. (Wilke et al., 1999) IDPs fluctuate depending on body position and can increase abruptly due to activities such as weight lifting, jumping/landing, and Valsalva maneuvers, resulting in radially-directed impact loading on the AF. (Gatt et al., 1997; Mustafy et al., 2016, 2014; van Loon et al., 2003) Abrupt spinal loading, in the presence or absence of IVD degeneration (IVDD) can induce damage resulting in IVD herniation (IVDH) of the NP through the development of focal defects in the AF (**Figure 1**). (Iencean, 2000) Patients with IVDH often undergo discectomy to remove the herniated IVD fragments; however, these remnant focal defects are not surgically repaired. Instead, the outer AF is left to heal on its own through a process that generates fibrotic scarring. (Torre et al., 2018) This scarring does not restore the microarchitecture or mechanical integrity of the AF which is evidenced by re-herniation rates of 9-25% at four and ten years follow-up in discectomy patients. (Atlas et al., 2005; Guterl et al., 2013; Hu et al., 1997) Furthermore, surgical intervention can result in focal AF injury. More specifically, patients with early-to moderate IVDD (an aberrant cell-mediated inflammatory process initiating in the NP) may benefit from early-stage interventions including NP replacement (NPR). However, pre-formed NPRs require injury to the AF to implant the device in the center of the IVD (**Figure 1**). (Yue et al., 2008) Thus, following implantation the AF needs to be closed to prevent subsequent herniation. Taken together, a need exists to develop a mechanically competent AF repair biomaterial, which allows for the immediate closure of focal AF defects and mimics the native structure and function of the outer AF to guide regeneration of native tissue. (Sharifi et al., 2014)

**Figure 1:**
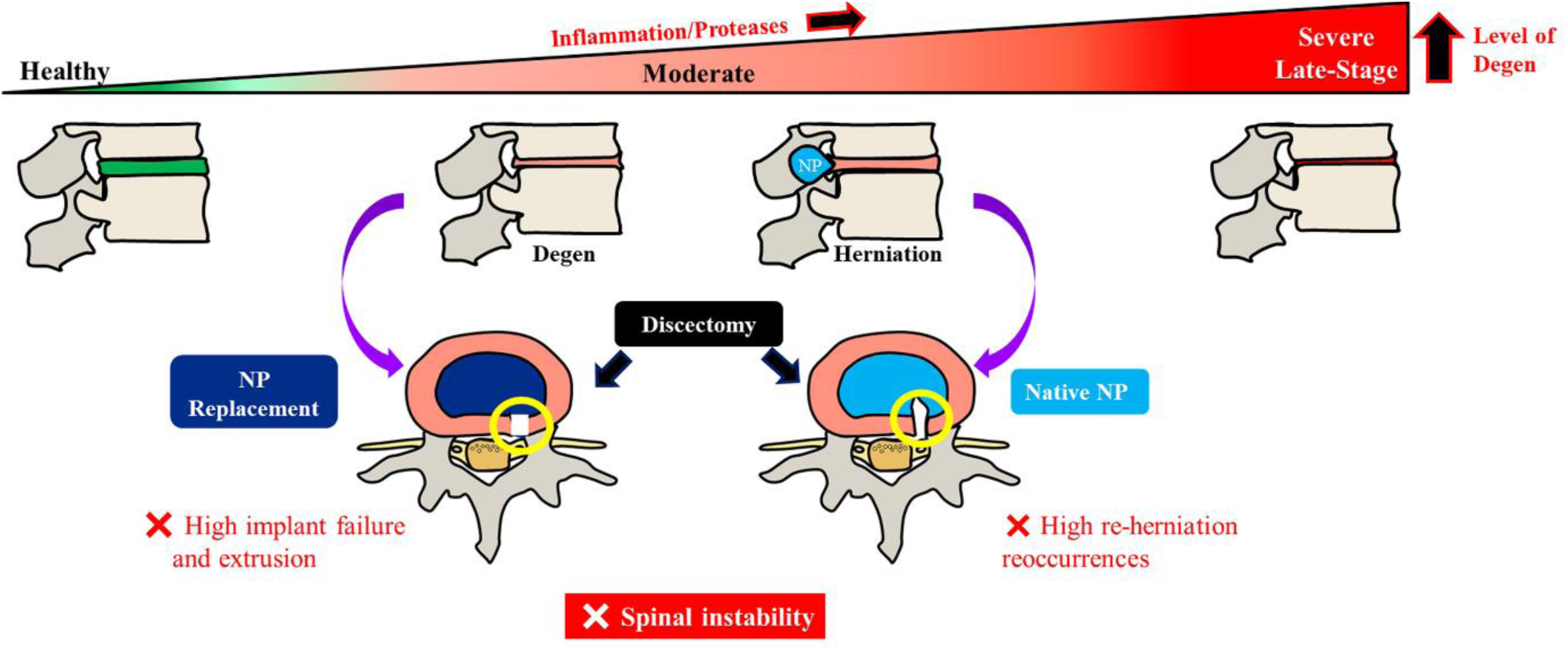
Representative image depicting the disc degenerative cascade. Increase in inflammation and proteases lead to increases in disc degeneration. IVDs suffering from moderate disc degeneration and herniation which are surgically repaired via discectomy result in a remnant annular defect (yellow circle). This allows for a pathway of least resistance for NP tissue to extrude/herniate; thus, leading to spinal instability.

In response to this need, there have been several attempts towards developing an outer AF closure system, including sutures (e.g., X-Close; Annulex) (Bailey et al., 2013; Chiang et al., 2012), adhesives (e.g., cyanoacrylates, glutaraldehyde-albumin, fibrin glues) (Guterl et al., 2014; Kang et al., 2017), or obstructive devices (e.g., Barricaid) (Lange et al., 2018; Wilke et al., 2013). However, these approaches do not possess the same composition, microarchitecture, or mechanical properties of the native AF, which has been shown to play a crucial role in AF function. (Iatridis et al., 2013) Thus, over the last decade increasing advancements in creating tissue-engineered scaffolds for the repair and regeneration of AF focal defects have been investigated. Many groups have focused on developing void-filling biomaterials that can replace missing AF tissue throughout the full-thickness (~5-6mm depth) of the AF. These include genipin-crosslinked fibrin hydrogels (Cruz et al., 2018; Frauchiger et al., 2018; R. Long et al., 2016), riboflavin cross-linked collagen gels (Pennicooke et al., 2018; Sloan et al., 2017), disc-like angle ply structures (DAPS) (Martin et al., 2017, 2014; Nerurkar et al., 2010), and synthetic poly(trimethylene carbonate) scaffolds (Pirvu et al., 2015). These innovative biomaterials have previously demonstrated the ability to support cell adhesion, proliferation, and ECM production, and are currently being evaluated for their mechanical competency and ability to assist in the restoration of functional spinal unit (FSU) kinematics. Of significant note, full-thickness AF repair biomaterials undergoing kinematic testing were sequestered within the IVD using an auxiliary synthetic membrane or ‘patch’ secured to the outer AF. (R. Long et al., 2016; Pirvu et al., 2015) Taken together, this suggests that a combinational approach may be required for successful IVD repair by utilizing an outer AF closure that can 1) resist impact loading, 2) aid in restoring FSU kinematics, and 3) secure a full-thickness AF repair biomaterial.

Thus, our lab has developed a biological scaffold, which is a biomimetic outer annulus fibrosus repair patch (AFRP) comprised of multiple sheets of type I collagen. AFRP’s are fabricated from decellularized porcine pericardium with its innate collagen fibers aligned with a ±30° “angle-ply” microarchitecture mimicking the native AF. (McGuire et al., 2016) Additionally, the AFRPs mechanical properties have been previously characterized demonstrating similarity to human AF; however, its ability to withstand high-impact loading warranted enhancement. (Borem et al., 2017) The AF’s ability to resist radially-directed impact is thought to be supported in part by the energy dissipative function of GAG found within the ILM. (Nerurkar et al., 2011; Nerurkar et al., 2011; Perie et al., 2006) Thus, we hypothesized the impact resistance of the AFRP would be increased via the inclusion of a GAG-based hydrogel ILM. Additionally, when using a combinational approach for AF repair, which includes both an outer AF closure device and a full-thickness AF repair biomaterial, it is essential to understand the contribution of each to FSU kinematics. Thus, the objectives of this research were to: 1) characterize the radially-directed impact resistance of the AFRP following incorporation of a GAG-based ILM, 2) determine the relative contributions of an AFRP and AF void-filler towards restoring injured FSU axial and torsional kinematics, and 3) evaluate the AFRP’s ability to provide adequate closure of an annular focal defect by preventing native IVD tissue herniation.

## 2. Materials and Methods

### 2.1 FABRICATION OF ANNULUS FIBROSUS REPAIR PATCHES (AFRPs)

Multi-laminate “angle-ply” AFRPs were developed and assembled from decellularized porcine pericardium to form tri-layer scaffolds and crosslinked in 6mM 1-ethyl-3-(3-dimethylaminopropyl) carbodiimide hydrochloride (EDC) with 1.2mM N-hydroxysuccinimide (NHS) as previously described. (Borem et al., 2017; McGuire et al., 2016) AFRPs were maintained in a phosphate buffered saline (PBS) storage solution containing protease inhibitor at 4°C for up to two weeks prior to testing.

### 2.2 HA-GEL PREPARATION AND AFRP ENRICHMENT

Hyaluronic acid (HA)-gel was prepared by reconstituting 1% sodium hyaluronate (a water-soluble salt form of HA) in PBS. AFRPs were then enriched by injecting the HA-gel between the AFRP lamellae (~62.5ug/mg) to form a GAG-based inter-lamellar layer (**Figure 2A&B**).

**Figure 2:**
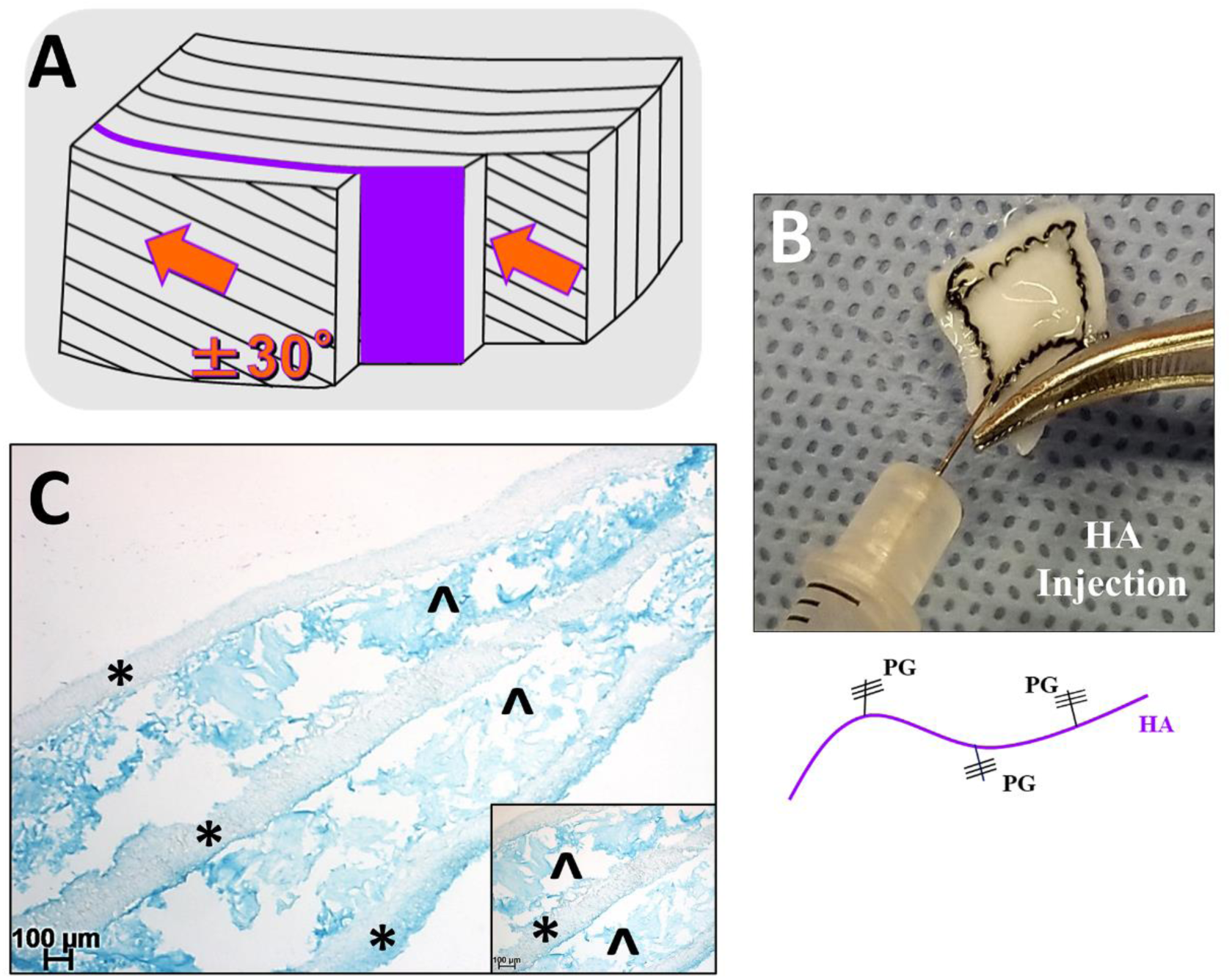
Glycosaminoglycan (GAG) enrichment of the AFRP. **A)** Representative schematic illustrating the proteoglycan-rich interlamellar matrix between native AF lamellae. **B)** Injection of hyaluronic acid (HA-non-sulphated GAG) between the AFRP lamellae. **C)** Representative image depicting GAG-enriched AFRP [**\*** indicates AFRP layer; ^ indicates positively stained GAG] (insert: magnified image of GAG-enriched AFRP).

### 2.3 HISTOLOGICAL STAINING FOR CONFIRMATION OF INTER-LAMELLAR GAG

AFRPs were fixed in 10% neutral-buffered formalin for 24 hours before undergoing successive washes in graded ethanol, xylene, and paraffin followed by embedding and sectioning to 5µm thickness. Slides were stained with Alcian Blue (1% Alcian Blue in 3% aqueous acetic acid; pH 2.5) and counterstained with 0.1% aqueous Nuclear Fast Red for visualization of glycosaminoglycan deposition. Histological images were then captured on a Zeiss Axio Vert.A1 microscope with AxioVision software (SE64 Rel. 4.9.1 SP08-2013).

### 2.4 IMPACT RESISTANCE TESTING

Radial impact strength testing was performed according to methods previously described. (Borem et al., 2017) Briefly, representative samples of crosslinked and crosslinked + HA-gel enriched AFRPs (3-ply; n=15/group; AFRP dimensions: 12 mm (L) × 12 mm (W) × 0.75mm (T)) were tested using a custom designed free-fall impact testing drop-tower. The base platform of the drop-tower consisted of a tissue-holding clamp and four vertical rails, which guided a free-falling platform. The tissue holding apparatus consisted of two stacked blocks lined with coarse-grit sandpaper each having an aligned thru-hole of 6.25mmØ. AFRPs were interposed between the two blocks centered over the two thru-holes. Subsequently, a 6mmØ steel ball attached to a 3-inch pushrod was placed in contact with the AFRP via the thru-hole in the superior block. Various weights ranging from 0.32-0.65kg were stacked on the free-fall platform, which was then dropped from a constant height of 0.254m. Impact energy (E) was calculated using the equation for kinetic energy, 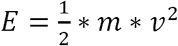, where m = mass and v = velocity. The resultant ball-burst pressure was calculated given the maximum force at rupture and its relationship with ball-burst pressure and geometric constraints according to established procedures.

### 2.5 PREPARATION OF FUNCTIONAL SPINAL UNITS

Bovine caudal tails, from 2-3-year-old calves, were obtained from a local abattoir and transported on wet ice to the lab within an hour. Excess tissue and ligaments surrounding the vertebral bodies and intervertebral discs were removed via dissection, and functional spinal units (FSUs: vertebrae-IVD-vertebrae) with posterior elements attached were isolated via a bandsaw. FSUs were harvested from two caudal levels (cc1-2 to cc2-3) and were potted using 3mm steel rods and urethane potting resin to prevent slippage of the samples during testing. Prior to testing, FSUs were wrapped in gauze saturated with storage solution and stored at −80°C. Samples were thawed within the sealed zip-lock bag, which was submerged for four hours in PBS at ambient temperature; thus, to prevent excessive tissue swelling.

### 2.6 BIOMECHANICAL EVALUATION OF FUNCTIONAL SPINAL UNIT AXIAL AND TORSIONAL KINEMATICS

FSU’s (n=5/group) underwent kinematic testing according to methods previously described by our group, with modification. (Borem et al., 2017) Briefly, FSU’s (n=5/group) were tested sequentially in the following groups: Intact, Repair with only an HA-gel enriched AFRP, and Repair with HA-gel enriched AFRP + AF tissue plug (**Figure 3A**). To initiate injury of the FSU, an annulotomy was performed by perforation of the IVD using a 6mmØ biopsy punch (7mm depth) and subsequent removal of the AF tissue. The AFRP repair only group consisted of the closure of the outer annular defect with an AFRP (AFRP dimensions: 7mm (L) × 7mm (W) × 0.75mm (T)), which was secured via a 4-0 FiberWire suture. Suturing of the AFRP consisted of suturing at the four corners with sutures being passed through the AFRP 1-2mm from the edge of the patch. Suture throws were made in alignment with the ±30° collagen fibrils of the AFRP. The AFRP + AF tissue plug group consisted of the restoration of the full-thickness annular defect using the previously incised native AF tissue recovered during the annulotomy, and the tissue was secured in place with an AFRP as described above.

**Figure 3:**
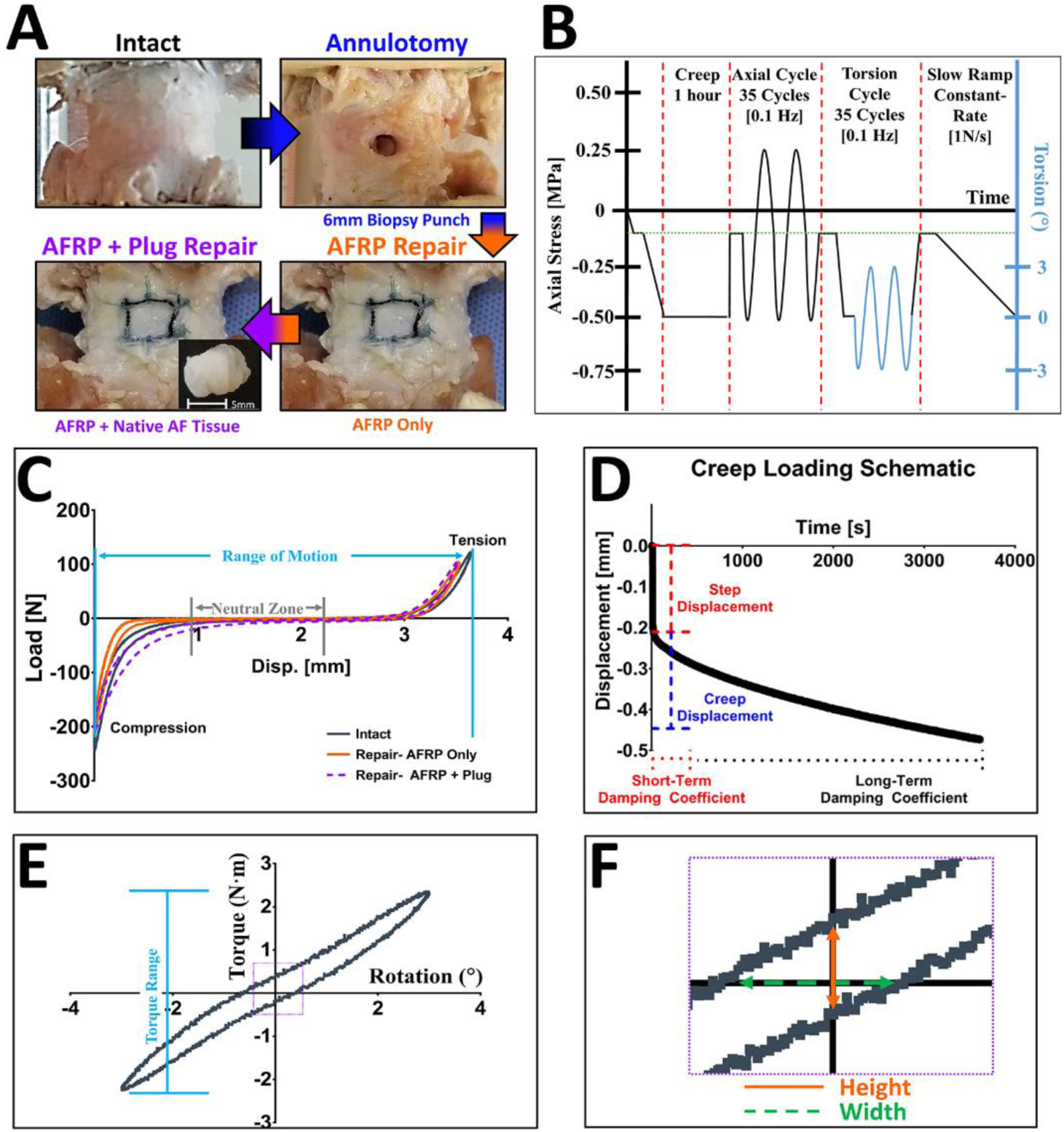
Study design for *in situ* kinematic testing of bovine IVD functional spinal units (FSUs) using GAG-enriched AFRPs. **A)** Representative images of the progression of testing groups (plus pictorial image of annular injury) and preparation for kinematic testing of FSUs. **B)** Loading scheme for FSU testing depicting creep loading, axial cyclic tension-compression loading, torsional loading, and slow constant-rate ramp loading. **C-E.1)** Representative graphs depicting the axial, creep, and torsional response of an IVD and its associated parameter measures. **E.2)** Graph depicting the torsional hysteresis height and width; parameters were calculated based on the dimensions of their respective geometric axes (orange solid line = height; green dotted line = width).

FSU’s first underwent creep loading on a Bose ElectroForce (model: 3220, TA Instruments) equipped with a 100-lb. load cell and a test chamber filled with 1xPBS/protease inhibitor at 25°C. Samples were loaded to a mean amplitude level of −0.125 MPa and then underwent a 1-hr. creep period at −0.50 MPa. Samples were then transferred to a servohydraulic test frame (model: 8874, Instron) fitted with a 20kN load cell, and subjected to 35 cycles of axial compression (−0.50 MPa) and tension (0.25 MPa) at 0.1 Hz. Compression was then maintained at −0.50 MPa as samples underwent 35 torsional cycles to ±3°. Finally, samples underwent a slow-rate compressive ramp (1 N/s) from a mean amplitude level of −0.125 MPa to −0.50 MPa (**Figure 3B**). Tensile and compressive stiffness was determined using a linear fit of the loading force-displacement curve from 60-100% of the 35^th^ cycle (**Figure 3C**). The axial range of motion was defined as the total peak-to-peak displacement of the IVD. Axial neutral zone (NZ) length was determined by fitting a third-order polynomial to the data and finding the maxima and minima with the correlating range between the peaks. A non-linear constitutive model was fit to the creep data (**Figure 3D**) using GraphPad Prism 7 software to yield elastic (η) and viscous (*·*) damping coefficients for the short-term (*η*_1_and Ψ_1_) and long-term (*η*_2_ and Ψ_2_), as described previously. Torsional stiffness was calculated from a linear fit of the loading torque-rotation curve of the 35th cycle. Torque range and axial range of motion (RoM) was calculated as the peak-to-peak torque and displacement, respectively. Changes in torque hysteresis were calculated by measuring the height and width of the curve along the geometric axes as shown in **Figure 3E&F**. The constant-rate slow-ramp compression stiffness was determined using a linear fit of the slow-ramp load-displacement response.

### 2.7 Compression to failure of Repaired FSUs

Following kinematic testing of FSUs, repaired samples were loaded to a mean amplitude level (−0.125 MPa) and then compressed at a rate of 300 mm/min until FSU failure (i.e., contact of the adjacent vertebral endplates and/or AFRP failure/AF Plug herniation). Ultimate compressive strength was calculated from the sample load and the initial cross-sectional area of the IVDs. Digital video was used to record mode and time of failure.

### 2.8 Statistics

Statistical analysis was performed on raw data using GraphPad Prism 7 software. Results are represented as mean ± standard error of the mean (SEM), and significance was defined as (p≤0.05). Impact resistance data was evaluated using a one-way ANOVA with Dunnett’s post-hoc analysis. Kinematic data was evaluated using a one-way repeated measures ANOVA followed by Dunnett’s post-hoc analysis.

## Results

### 3.1 Confirmation of GAG-enrichment of AFRPs

HA-gel enriched AFRPs were histologically evaluated to confirm the presence of GAG between the AFRP lamellae following injection. Positive Alcian blue staining of the crosslinked + HA-gel AFRPs demonstrated positive staining within the inter-lamellar region confirming the presence of the GAG (**Figure 2C**).

### 3.2 Impact testing

Multi-laminate AFRPs underwent radially-directed impact loading to evaluate its resistance to failure during the application of sudden changes in IDP (**Figure 4A&B**). Mean impact strength of non-crosslinked, crosslinked, and crosslinked + HA-gel AFRPs was 1.51 ± 0.06 MPa, 2.04 ± 0.03 MPa, 2.57 ± 0.02 MPa, respectively (**Figure 4C**). Additionally, the average kinetic energy absorbed by non-crosslinked, crosslinked, and crosslinked + HA-gel AFRPs was 0.51 ± 0.02 J, 0.69 ± 0.01 J, 0.87 ± 0.01 J, respectively (**Figure 4D**). Expectedly, impact strength was statistically increased following crosslinking (p<0.001) and crosslinking + HA-gel (p<0.001) compared to non-crosslinked AFRPs. Furthermore, impact energy was statistically increased in crosslinked AFRPs (p<0.001) and crosslinked + HA-gel AFRPs (p<0.001) compared to non-crosslinked AFRPs.

**Figure 4:**
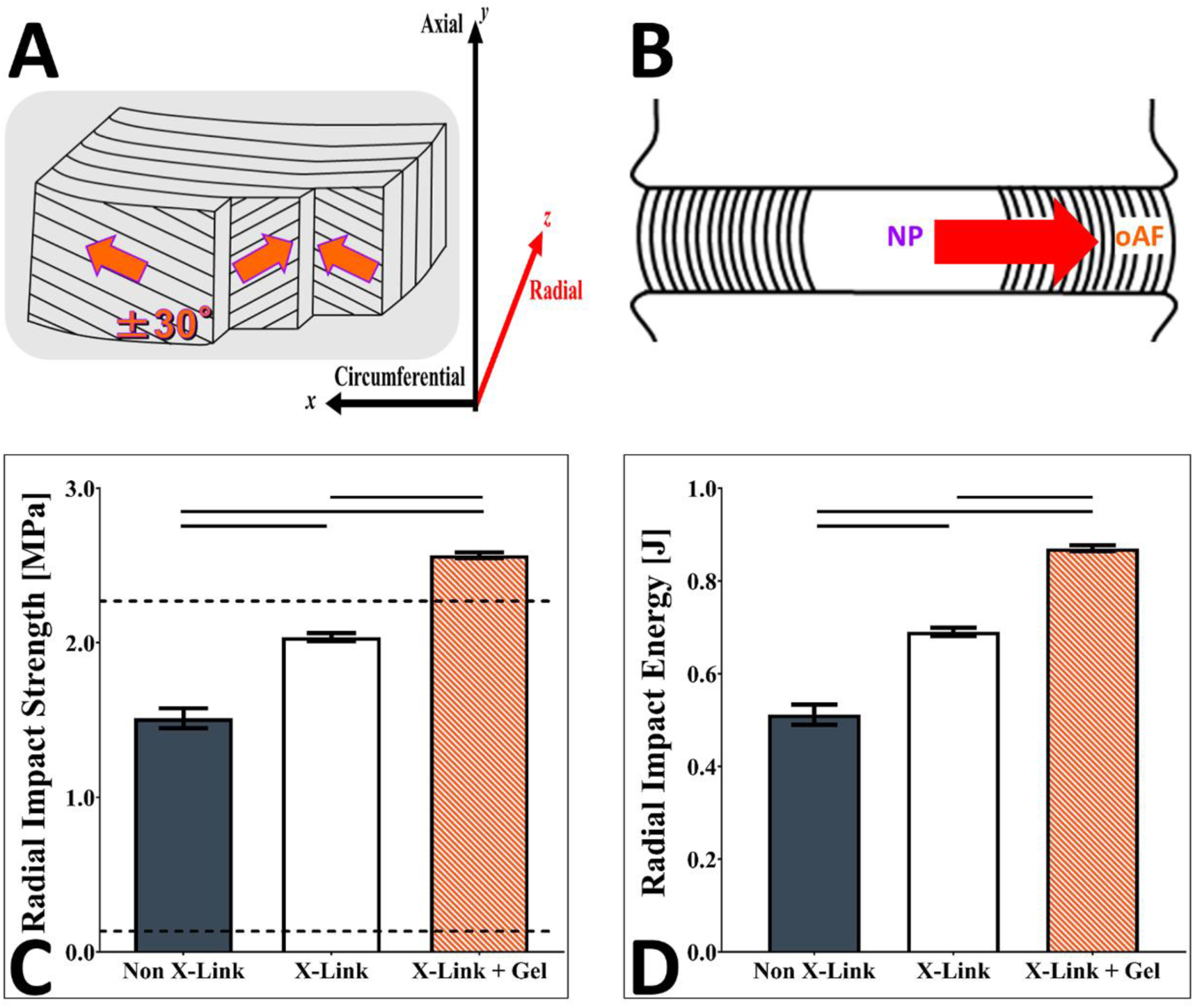
Biomechanical evaluations of non-crosslinked, crosslinked, and crosslinked + HA-gel AFRPs subjected to radial impact burst strength. **A&B)** Representative schematics illustrating the radial loading axis the NP exerts on the AF through increases in intradiscal pressures (IDPs). **C)** Graph depicting the average radial impact strength withstood by AFRP testing groups. Dotted line indicates reported human values of IDPs 0.1-2.3 MPa. **D)** Graph depicting the average radial impact kinetic energy absorbed by the AFRP testing groups.

### 3.3. Kinematic Results

Axial and torsional FSU kinematics were evaluated to determine the impact of AF focal defect repair (created via annulotomy) between the outer AFRP scaffold and combinational approach using a full-thickness AF tissue plug (excised intact AF tissue). Of note, completion of kinematic testing with the AF tissue plug could not be successfully achieved without sequestering it within the IVD with an AFRP. This was due to the pressurization of the native NP, which caused the AF tissue plug to extrude/herniate prior to mechanical loading (**Figure 5**).

**Figure 5:**
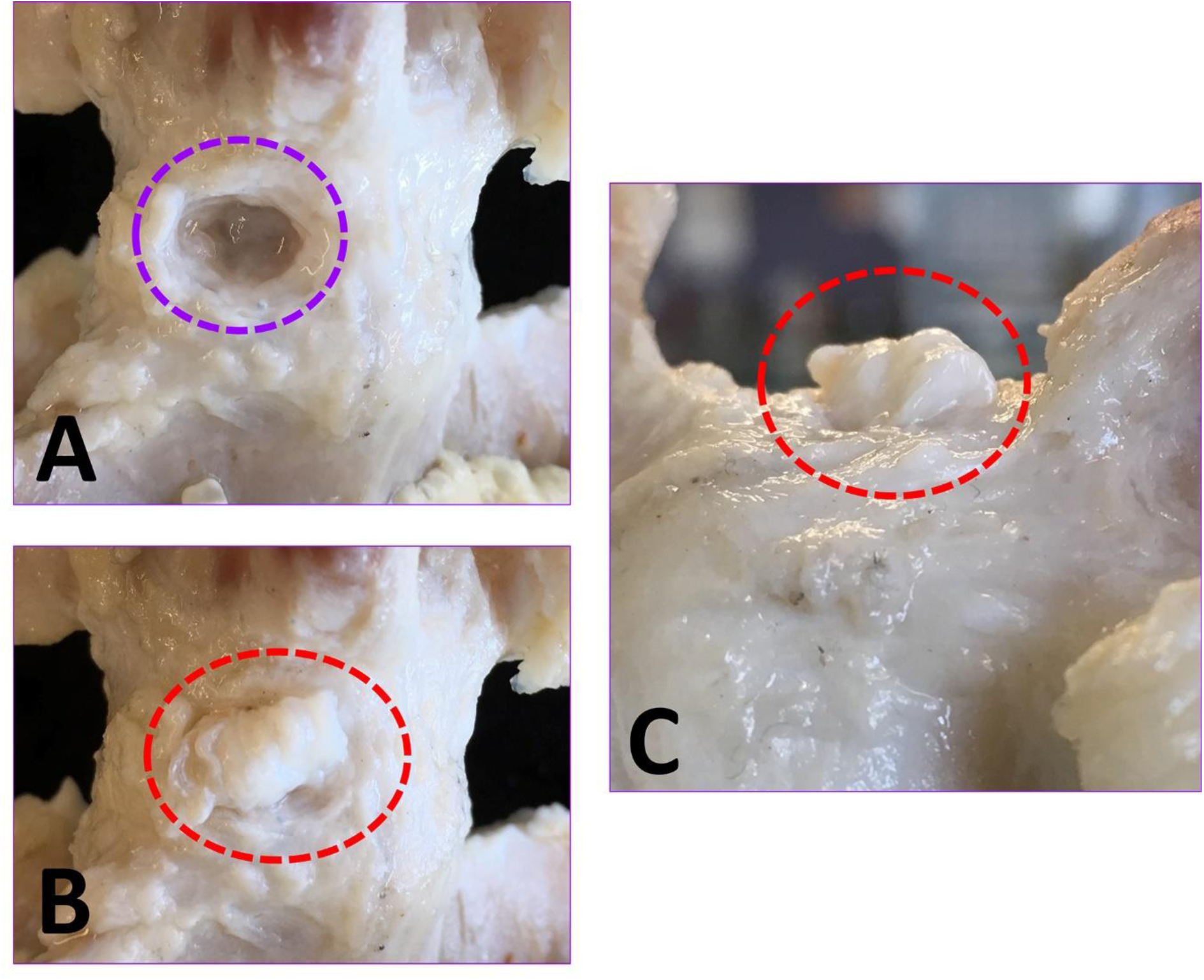
Pressurization of the IVD prevented repair with the native AF Plug only without an AFRP. **A)** Annular defect following annulotomy procedure (purple dotted circle). **B&C)** Top and Side views of an AF Plug placed into FSU illustrating the immediate extrusion out of the defect following implantation (AF Plug centered in red dotted circle).

#### 3.3.1 Axial Cyclic Loading

Axial cyclic kinematic testing of FSU’s repaired with the AFRP scaffold (with and without AF tissue plug) did not significantly alter axial compressive stiffness, slow ramp compressive stiffness, or tensile stiffness kinematic parameters compared to intact controls (**Figure 6A&B**). Conversely, repair of FSUs with only the AFRP scaffold resulted in a statistical increase compared to intact controls in axial range of motion (AFRP: 4.03 ± 0.27 mm, Intact: 3.45 ± 0.29 mm; p=0.045) and neutral zone (AFRP: 1.55 ± 0.11 mm, Intact: 1.28 ± 0.13 mm; p=0.037) (**Figure 6C&D**). However, incorporation of the native AF tissue plug, sequestered by an AFRP, restored these parameters to intact values (axial range of motion: 3.67 ± 0.29 mm; p=0.476 and neutral zone: 1.31 ± 0.11 mm; p=0.908).

**Figure 6:**
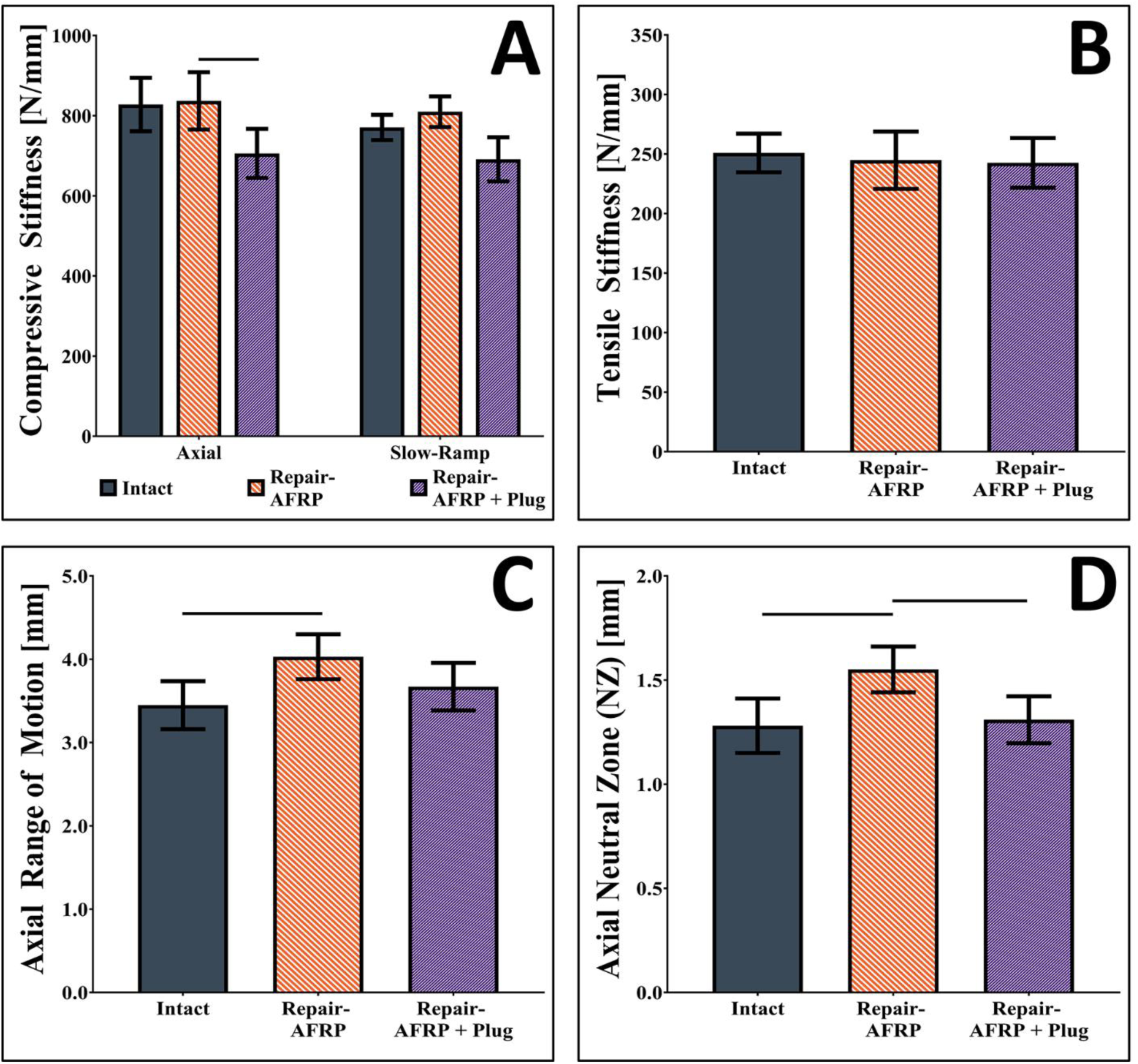
Axial cyclic tension-compression kinematic testing results of bovine IVD FSUs. Graphs depicting **A)** axial and slow ramp compressive stiffness, **B)** tensile stiffness, **C)** axial range of motion, **D)** axial neutral zone. Solid lines connecting groups indicates significant difference (p<0.05) compared to uninjured controls.

#### 3.3.2 Creep loading

Creep loading of FSU’s repaired with the AFRP scaffold (with and without AF tissue plug) did not significantly alter the step displacement or short-term elastic damping coefficient (**Figure 7**). However, repair of FSUs with only the AFRP scaffold resulted in a statistical increase compared to intact controls in creep displacement (AFRP: 1.29 ± 0.07 mm, Intact: 1.56 ± 0.07 mm; p=0.03), long-term elastic damping coefficient (AFRP: Ψ_2_:201.30 ± 14.97 N/mm, Intact: 140.10 ± 3.53 N/mm; p=0.022), short-term and long-term viscous damping coefficients (AFRP: η_1_: 7532.40 ± 1251.02 Ns/mm and η_2_:46.59 × 10^4^± 3.29 × 10^4^Ns/mm, respectively, Intact: η_1_:4.73 × 10^3^ ± 1.07 × 10^3^ Ns/mm and η_2_:3.70 × 10^5^ ± 0.11 × 10^5^ Ns/mm, respectively; p_1_=0.003 and p_2_=0.024). However, IVD repair incorporating the native AF tissue plug, sequestered by an AFRP, restored these parameters to intact values for creep displacement (1.74 ± 0.06 mm; p=0.134), long-term elastic damping coefficient (Ψ_2_:144.48 ± 7.50 N/mm; p=0.751), short- and long-term viscous damping coefficient (η_1_: 5839.60 ± 1327.04 Ns/mm and η_2_:31.33 × 10^4^± 1.92 × 10^4^ Ns/mm, respectively; p_1_=0.538 and p_2_=0.058).

**Figure 7:**
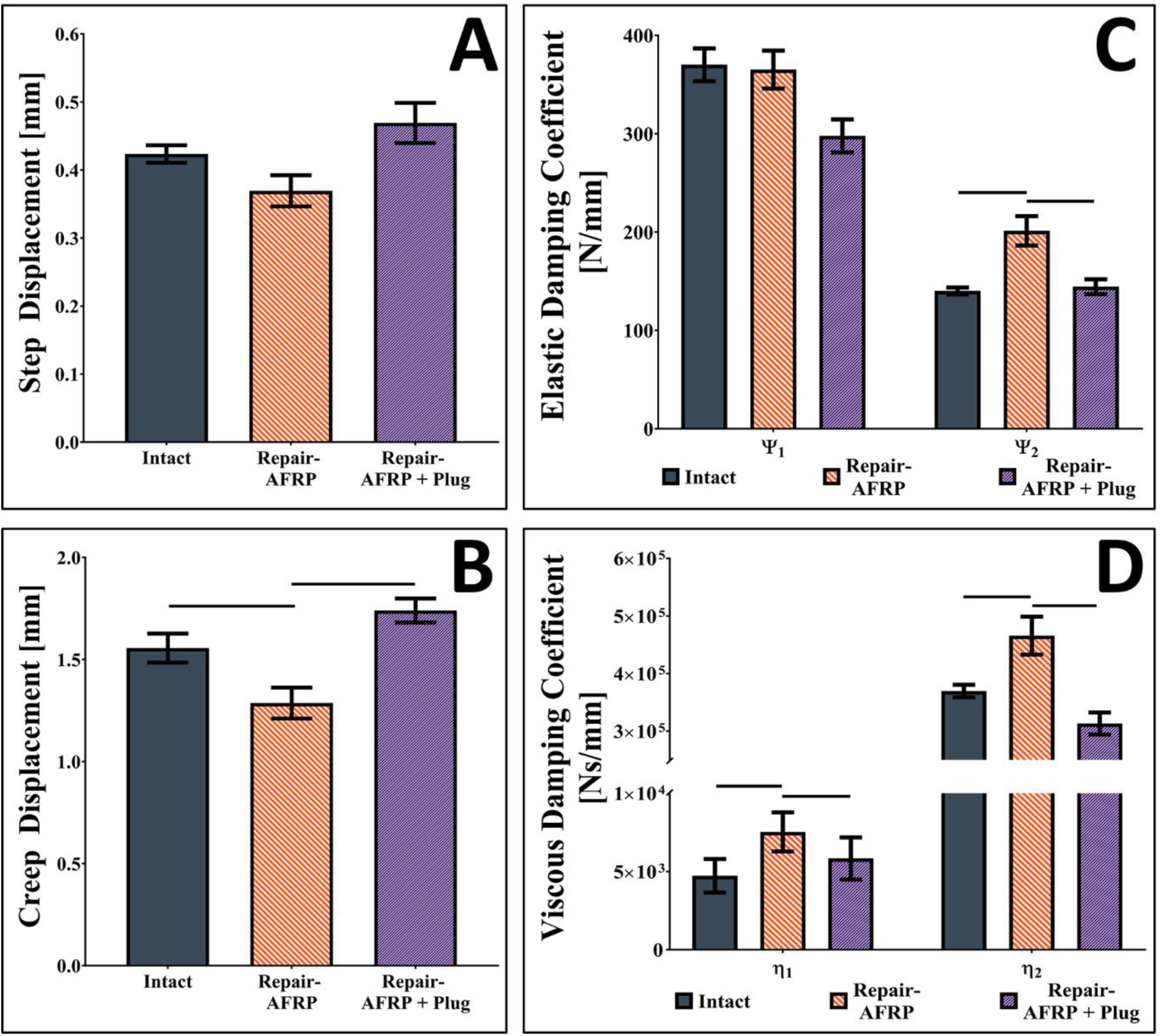
Creep kinematic testing results of bovine IVD FSUs. Graphs depicting **A)** step displacement, **B)** creep displacement, **C&D)** elastic & viscous damping coefficients. Solid lines connecting groups indicates significant difference (p<0.05) compared to uninjured controls.

#### 3.4.3 Torsional Cyclic Loading

Focal injury to the AF resulted in significant changes in FSU torsional kinematics, which were not restored by any repair methodology (**Figure 8**). Torsional rotation of FSU’s repaired with only the AFRP or in combination with the AF tissue plug demonstrated a significant decrease in torsional stiffness (0.48 ± 0.02 Nm/°: p=0.001 and 0.53 ± 0.03 Nm/°: p=0.024, respectively) (**Figure 8A**), torque range (3.45 ± 0.12 Nm: p=0.0002 and 3.68 ± 0.17 Nm: p=0.022, respectively) (**Figure 8B**), and torque hysteresis height (0.28 ± 0.03 Nm: p=0.028 and 0.30 ± 0.04 Nm: p=0.023, respectively) (**Figure 8C**) compared to intact FSUs (0.66 ± 0.02 Nm/°, 4.60 ± 0.10 Nm, 0.42 ± 0.05 Nm, respectively).

**Figure 8:**
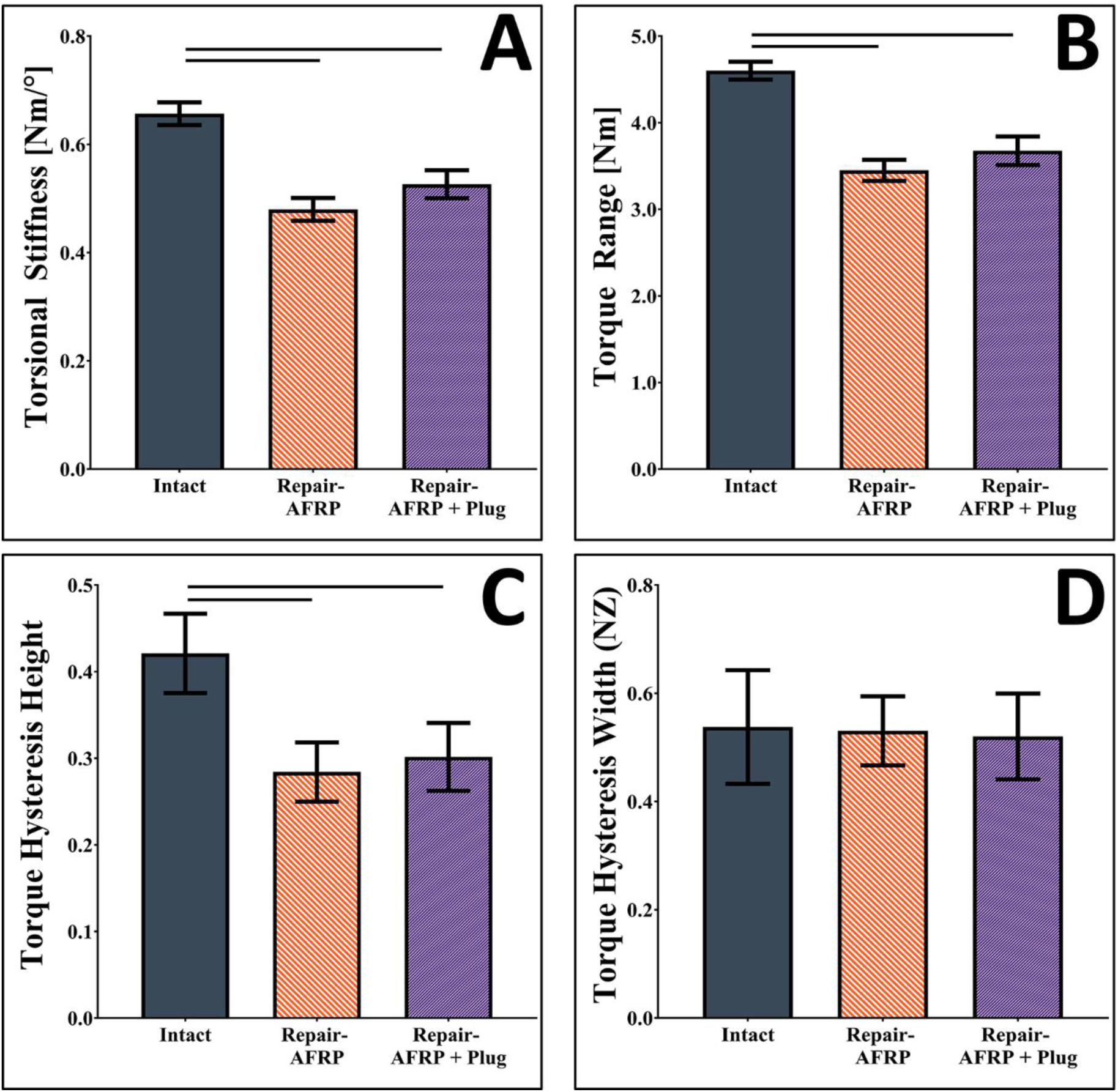
Torsional kinematic testing results of bovine IVD FSUs. Graphs depicting **A)** torsional stiffness, **B)** torque range, **C)** torque hysteresis width (*NZ*), **D)** torque hysteresis height. Solid lines connecting groups indicates significant difference (p<0.05) compared to uninjured controls.

#### 3.4.4. Compression to Failure

Axial compressive testing of FSU’s repaired with the AFRP scaffold (with and without AF tissue plug) were assessed for compressive failure strength concomitant with visual assessment. Testing demonstrated no failure in the AFRP attachment nor herniation of native IVD tissues. Testing was stopped once video recordings illustrated that vertebral endplates contacted each other, and an axial load was no longer applied. Furthermore, no statistical differences were observed between the repair groups ultimate compressive strength (AFRP: 5.28 ± 0.62 MPa; AF Plug: 3.84 ± 0.48 MPa; p=0.101) or compressive displacement (AFRP: 2.58 ± 0.25 mm; AF Plug: 2.96 ± 0.61 mm; p=0.619).

## Discussion

Focal injuries of the AF, produced by NP herniation or surgical trauma to enable implantation of an NPR, ultimately results in mechanical dysfunction of the IVD. (Guterl et al., 2013) Thus, repair of AF defects using mechanically competent implants that can withstand abrupt spinal loading and assist in restoring IVD kinematics are required. Herein, we characterized a multi-laminate AFRP scaffold containing a GAG-based ILM for its ability to resist radially-directed impact loading and its contribution to FSU kinematics when used in conjunction with a full-thickness AF tissue plug. The primary findings from these studies include: 1) incorporation of an ILM significantly increased the impact strength of the AFRP, 2) the AFRP enabled the restoration of axial FSU kinematics, and 3) the AFRP prevented herniation of native NP and a full-thickness AF tissue plug under applied physiological loading. Together, these results suggest that the AFRP can be used as a mechanical closure system to sequester NP and AF tissues and/or repair biomaterials within the IVD to assist in restoring its native mechanical function.

The first major finding from this study was that inclusion of an ILM composed of an HA-hydrogel between the layers of the AFRP scaffold resulted in a significant increase in its radial impact strength. To the best of the author’s knowledge, this is one of the first attempts to incorporate a GAG-based ILM within a biologic annulus repair scaffold to better recapitulate the anatomy and physiology of the native AF. GAGs, including chondroitin sulfate and HA, make up approximately 10-20% of the overall dry weight of the AF. (Perie et al., 2006; Roughley, 2004) Moreover, these molecules have been shown to play a functional role in contributing to the overall compressive properties of the IVD (e.g., compressive equilibrium modulus and peak stress) and helps the AF to resist IDP and prevent delamination and shearing of the AF lamellae. (Adam et al., 2015; Isaacs et al., 2014; Kirking et al., 2014; Nandan Nerurkar et al., 2011; Perie et al., 2006) Herein, we demonstrate that the inclusion of a GAG-gel between the layers of the AFRP can play a significant role in dissipating radial-directed impact loads. This is somewhat intuitive given they biochemical properties of GAGs and the load-bearing role they play in many tissues including heart valves and cartilage. (Basalo et al., 2004; Lovekamp et al., 2006; Wilson et al., 2009) In fact, our results suggest that incorporation of a GAG-gel ILM increases the AFRPs radial impact resistance from 3 g’s (non-crosslinked AFRPs (Borem et al., 2017)) to a conservative 10 g’s of gravitational force (equivalent to the force exerted when falling on your buttocks backward into low office chair (Allen et al., 1994), and 3x the force experienced on an aggressive roller coaster (Bibel, 2007)), both of which are beyond the magnitudes expected for normal activities of daily living following IVD repair.

The second major finding from these studies demonstrated that repair of AF focal defects in injured FSUs required a combinational approach, using both an outer AF closure and a full-thickness repair, to restore axial kinematics to intact levels. Moreover, when only an AFRP repair was applied, FSU kinematics were detrimentally changed; however, it was visually confirmed these changes were attributed to the migration of the native NP tissue into the void space of the focal AF defect and were not directly attributed to the AFRP itself. Furthermore, this migrated NP tissue had to be forcibly pushed back into the center of the IVD to replace the full-thickness AF tissue plug for subsequent testing. This pressurization phenomenon made it impossible for completing the full-thickness repair without the use of an outer AF closure patch which has been observed in other studies. (R. Long et al., 2016; Pirvu et al., 2015)

Notably, torsional kinematic parameters were not restored with either repair methodology. This could be contributed to the lack of integration with the surrounding annulus tissue. While the native full-thickness AF tissue plug occupied the structural void, lack of adhesion to surrounding tissue resulted in a lack of continuity. Therefore, it is presumed the use of a biocompatible adhesive could restore the continuity between the full-thickness AF tissue plug and the adjacent native annular tissue. However, previous investigations of AF repair biomaterials using adhesives have demonstrated only partial restoration of torsional kinematic parameters toward intact levels. (Likhitpanichkul et al., 2014; R. Long et al., 2016) Thus, the recovery of mechanical torsional properties may also be dependent on the restoration of circumferential residual stresses and pre-strains found within the AF. (Mengoni et al., 2017; Michalek et al., 2012) Accordingly, future investigations of AF repair biomaterials should take into consideration the restoration of native residual stresses and pre-strains to further assist in restoring torsional kinematic parameters.

The third major finding was that repair of the outer AF with the AFRP scaffold prevented herniation of both native NP and a full-thickness AF tissue plug repair. This is significant given the proposed role of this biological scaffold is to sequester either native tissue or implants which restore the native function of the IVD. In general, IVD function is dependent on the interaction of the NP and AF. More specifically, under compressive loading, the NP becomes pressurized and exerts radially-directed compressive forces on the AF that in turn help to reinforce the AF and prevents inward buckling of the lamellae. (Adam et al., 2015) Thus, the AFRP must be able to withstand these forces generated by the native tissue or by implant surrogates. Previous studies of AF repair biomaterials (with and without synthetic membranes) have demonstrated that under axial compressive loading these biomaterials experience the inclination to herniate at or below physiological stresses. (Cruz et al., 2018; R. Long et al., 2016; Long et al., 2017) Additionally, while other studies using similar AF repair biomaterials have demonstrated the ability to resist expulsion from the AF focal defect, a substantial portion of native NP tissue is often removed which in turn may presumably reduce the IVD pressurization. (Cruz et al., 2018; Likhitpanichkul et al., 2014) Therefore, these AF repair biomaterials may not be exposed to the full magnitude of radially-directed loading following the removal of native NP tissue. Therefore, under a more rigorous testing scenario (full pressurization of native IVD tissue), we have shown that the AFRP can resist the coordinate pressurization and radially-directed translation of native NP and a full-thickness AF tissue plug under physiological relative loads. Additionally, while macroscopic and video analysis of the AFRP illustrated minimal outward bulging during axial compressive loading, the AFRP showed no failure at the patch/suture interface nor did herniation/extrusion of the native NP or full-thickness AF tissue plug occur.

## Conclusions

In summary, AFRP’s demonstrated its ability to resist physiological impact loading IDPs, prevent herniation of a full thickness AF void-filler, and assisted in the restoration of spinal kinematic parameters. Moreover, kinematic testing results expressed herein demonstrated the necessity of both 1) a full thickness AF void-filler and 2) an outer annular closure biomaterial to restore spinal kinematic parameters to intact values. Therefore, to restore physiological functionality, surgical repair of IVD disorders should include both a full thickness annular replacement to fill the mechanical void and an AFRP biomaterial to reinforce the immediate closure of the AF.

### Limitations

As with any study, the authors acknowledge some limitations within this study. Concerning impact testing, the authors recognize that the ILM is composed of a variety of components, all of which play a crucial role in the maintenance of physiological function of the native AF. Although, the purpose of this study was to improve the impact resistance of AFRPs, and the proteoglycan-component of the ILM expresses a viscous response that was hypothesized to serve as a shock absorber to dissipate the force applied during impact loading. An additional limitation acknowledged by the authors, regarding impact testing, is that AFRPs were not evaluated over multiple cycles for their impact fatigue strength. However, the magnitudes of impact pressures used during testing are considered infrequent (e.g., falling on your buttocks, riding a rollercoaster, and car accidents), and not expected to be a common occurrence for people who recently undergone IVD repair. Additionally, for this study, AFRPs impact strength was evaluated for its ability to resist the complete absorption of the impact energy using a 6mmØ steel-ball to represent a large annular defect (Guterl et al., 2013); however, this is thought to serve as a “worse-case” scenario, as the inclusion of a full-thickness AF repair biomaterial will also absorb IDP exerted by the NP and dissipate the energy applied to the AFRP. Concerning kinematic testing, one limitation was FSUs were not re-equilibrated for 8 hours submerged in saline between testing cycles, which has been recently shown by others to effect FSUs kinematics. (Bezci and O’Connell, 2018) Lastly, the authors acknowledge that laboratory-based mechanical testing performed herein may not directly reflect the potential changes of accumulation in damage over time when implanted *in vivo*.

## Acknowledgments

Research support for the Ortho-X lab has been provided in part by the National Institute of General Medical Sciences of the National Institutes of Health [award number: 5P20GM103444-07]. RB is supported by the National Science Foundation Graduate Research Fellowship [grant number: 2011382]. Additionally, we would like to thank Mr. Joshua Walters for his assistance during the kinematic potting process and review of the manuscript.

